# The Impact of Age and Sex on Cerebral and Large Artery Stiffness and the Response to Pulse Pressure

**DOI:** 10.1101/2025.04.01.646687

**Authors:** Young D Choi, Prabhuli Kapadia, Jamie Spiegel, Abigail E Cullen, Jessica LaFarga, Ashley E Walker

## Abstract

Vascular aging is characterized by a tandem increase in pulse pressure and large elastic artery stiffness. Greater stiffness of the large arteries leads to elevated pulse pressure transmitted into the cerebral circulation, causing dysfunction. However, little is known in females about age-related stiffening of the arteries and the impacts of high pulse pressure on the cerebral vasculature. To examine the effects of sex and age on the cerebral artery response to pulse pressure, we studied cerebral arteries collected from young and old female and male C57BL/6 mice. Isolated cerebral arteries were exposed *ex vivo* to static pressure, low pulse pressure, and high pulse pressure. Exposure to high pulse pressure impaired endothelium-dependent dilation in cerebral arteries from young female and male mice, with impairments also occurring in young female cerebral arteries after exposure to low pulse pressure. In contrast, exposure to low or high pulse pressure did not impact cerebral artery endothelium-dependent dilation for old male or female mice. During exposure to high pulse pressure, young females had higher cerebral artery compliance compared with young males and old females. Old mice also had higher cerebral artery passive stiffness and aortic pulse wave velocity compared with young mice. We also found age and sex differences in arterial wall thickness, collagen and elastin content, and matrix metalloproteinase 9 expression. Taken together, young female mice have more compliant cerebral arteries, which are more susceptible to endothelial dysfunction caused by pulse pressure.

## INTRODUCTION

Increases in arterial stiffness are primary features of the aging vasculature^1,2^ and are associated with neurodegenerative diseases and cognitive decline.^3,4,5^ The stiffening reduces the large arteries’ ability to dampen pulse pressure {PP}, resulting in undamped PP being transmitted down the vascular tree.^6^ The cerebral circulation may be particularly susceptible to increases in PP due to the high blood flow.^7-9^ Indeed, in humans, greater large artery stiffness and/or PP are associated with impaired cerebral blood flow and more white matter hyperintensities.^7,10,11^ Furthermore, previous work by us and others in rodent models demonstrates that greater large artery stiffness or high PP is associated with impaired cerebral blood flow, cerebral artery endothelial dysfunction, and a more permeable blood-brain barrier.^12-14^ However, there remain gaps in knowledge about the association of arterial stiffness and/or PP with cerebrovascular dysfunction, particularly the influence of sex differences and the role of cerebral artery stiffness.

Despite prominent cerebrovascular disease and Alzheimer’s disease {AD} risk in older females more is known about the mechanisms of dysfunction in males. Over 60% of AD patients are females^15^ and progression from mild cognitive impairment to AD advances at a faster rate in females compared to males.^16^ Differences in the mechanisms or consequences of large artery stiffening could drive these differences in disease risk. In females, there is a stronger association between mortality and large artery stiffness compared with males.^17^ The rate of increase in arterial stiffness and cerebral pulsatility with aging is higher in females than males, corresponding to an acceleration in changes during menopause.^18^ In animal models, studies have shown female mice develop stiffer arteries and have a lower elastin-to-collagen ratio later in life when compared with male mice.^18,19^ While these previous studies indicate that the age-related increases in large artery stiffness are different with sex, little is known about how this impacts the cerebral vasculature. It was previously shown that cerebral arteries from old male mice are less susceptible to the dysfunction caused by high pulse pressure than arteries from young male mice^20^; but this relationship was not explored in female mice. Therefore, we set out to determine the impact of sex on the dysfunction caused by elevated pulse pressure in both young and old mice.

While there is an extensive understanding of the stiffening of large arteries with advancing age, less is known about the age-related changes to cerebral artery stiffness. The extracellular matrix comprises the major load-bearing components of the aorta, comprised of elastin and collagens I, III, and IV.^21^ In the large arteries, previous human studies show that advancing age is associated with fragmentation of elastin and increased collagen content and cross-linking.^22,23^ Meanwhile, collagen degradation occurs simultaneously from pro-inflammatory signaling, the same signaling that degrades elastin and activates smooth muscle cell proliferation.^24-26^ Transforming growth factor {TGF}, platelet-derived growth factors {PDGF}, and matrix metalloproteinase-9 {MMP9} are pro-inflammatory factors that are associated with stiffening of arteries.^27-29^ Similar to large arteries, human and rodent studies demonstrate greater stiffness of the cerebral arteries and arterioles with advancing age.^20,30,31^ This greater stiffness is associated with a greater wall thickness and collagen content in cerebral arterioles from older mice.^30^ Yet, it is still unknown if the cerebral arteries follow similar age-related changes to the extracellular matrix and signaling as large arteries, and further sex differences in the changes to cerebral artery structure with aging are unknown.

The goal of this project was to examine the interaction of age and sex on cerebral artery structure and the endothelial dysfunction caused by high PP. We hypothesized that exposure to high PP would cause a greater impairment in endothelium-dependent dilation in cerebral arteries from old female mice compared to young female and male mice, and old male mice. As females experience a more rapid increase in large artery stiffness progression, we further hypothesized that cerebral arteries from old female mice would have greater elastin breakdown and collagen accumulation compared to young females and young and old males. To test these hypotheses, we collected cerebral arteries from young and old, female and male C57BL/6 mice and exposed the arteries to *ex vivo* static pressure, low PP, and high PP, followed by measurement of endothelium-dependent vasodilation. We also examined the stiffness of the aorta and cerebral arteries, as well as structure by immunofluorescence and initial investigation into signaling mediators by gene expression.

## EXPERIMENTAL METHODS

### Animals

Female and male C57BL/6 mice were obtained from the NIA Aging colony at Charles River and studied at 6-7 or 23-24 months of age {n= 10-15/group}. All mice were placed on a phytoestrogen-free diet {Envigo, Teklad Global Soy Protein-Free Extruded Rodent Diet, 2920X} with food and water provided *ad libitum* and housed in the animal care facility on a 12/12-hour light-dark cycle at 24°C. Mice were euthanized by exsanguination under isoflurane immediately before *ex vivo* vascular reactivity studies. All animal procedures conform with the Guide to the Care and Use of Laboratory Animals {8^th^ edition, revised 2011} and were approved by the Institutional Animal Care and Use Committee at the University of Oregon. Animal characteristics are displayed in Supplemental Table 1.

### Aortic Stiffness

Aortic stiffness was measured *in vivo* by pulse wave velocity {PWV}. Mice were anesthetized with isoflurane {2% isoflurane in 100% oxygen} and were placed on a 37°C temperature-controlled heating pad to maintain body temperature. To record ECG, electrodes were placed on distal portions of limbs. Two Doppler transducers were each placed on the aortic arch and abdominal aorta. The distance between the two transducers was recorded and the Indus Doppler Flow Velocity System {Webster, Texas} was used to register and analyze the data. The time of a pulse to travel from the aortic arch to the abdominal aorta was measured and divided by the distance between the transducers. Two researchers independently analyzed all files and the average was taken. Any PWV measurement with confidence intervals over 15% was excluded.

### Vascular Reactivity

Endothelium-dependent vasodilation was assessed *ex vivo* in isolated posterior cerebral arteries {PCAs}. Post euthanasia, the PCA was dissected from the brain and cannulated onto glass micropipette tips in a myograph chamber {Danish Myo Technology, Hinnerup, Denmark} filled with 145 mM NaCl, 4.7 mM KCl, 2 mM CaCl_2_, 1.17 mM MgSO_4_, 1.2 mM NaH_2_PO_4_, 5.0 mM glucose, 2.0 mM pyruvate, 0.02 mM EDTA, 3.0 mM MOPS buffer, 10 g/L BSA, 7.4 pH at 37°C. Arteries were pressurized to 50 mmHg and all arteries were preconstricted with phenylephrine {1-6 µM to obtain 20-40% preconstriction}. Changes in luminal diameter in response to increasing concentrations of endothelium-dependent dilator acetylcholine {ACh: 1×10^−9^ to 1×10^−4^ M} were determined. Endothelium-independent dilation was measured by the change in lumen diameter to sodium nitroprusside {SNP: 1×10^−10^ to 1×10^−4^ M}.

### *Ex vivo* Pulsatile Pressure Application

PCA endothelium-dependent dilation in response to pulse pressure {PP} was assessed by *ex vivo* pulsation exposure. To measure the effect of pulsation, arteries were pulsed at a frequency of 400 beats per minute at 50% duty cycles for 30 minutes each. The mean arterial pressure was set to be of similar pressure {62.5 mmHg} between conditions; the low PP condition was set to cycle between 50 and 75 mmHg and the high PP condition was set to cycle between 37.5 and 87.5 mmHg. Since the exact PP in the PCA is unknown, the pressures were estimated based on the carotid artery PP in a mouse model of higher large artery stiffness.^12^

### Arterial Distension and Stiffness with Pulsatile Pressure Application

The change in PCA lumen diameter was assessed following 30 minutes of pulsation via a 30-second, high-frame rate video recording with MyoView software {Danish Myo Technology, Hinnerup, Denmark}. VasoTracker Image Analyzer {open source} was used to measure and average frame-by-frame diameters across five sections of the artery. To exclude anomalies in the diameter tracking, the minimum and maximum diameters had to occur in greater than 10% of the total frames. Absolute distension was calculated as D_max_-D_min_ and percent distension as {D_max_-D_min_}/D_max_ ^*^ 100, where D_max_ is the maximum lumen diameter and D_min_ is the minimum lumen diameter.

β stiffness index was calculated as ln{P_high_/P_low_}/{{D_max_ D_min_}/D_min_}, where P_high_ is the highest pressure and P_low_ is the lowest pressure during the pulse application.^32^ Compliance was calculated as {A_max_-A_min_}/{P_high_/P_low_}, where A_max_ is the maximum lumen area and A_min_ is the minimum lumen area. Peterson modulus of elasticity {Ep} was calculated as D_min_ {P_high_ -P_low_}/{D_max_-D_min_}.^33^

### Passive Stiffness

After a 60-minute incubation in a calcium-free solution to eliminate the effect of myogenic tone, passive arterial stiffness was measured *ex vivo* in the PCAs. Medial wall thickness and lumen diameter were measured following increasing intraluminal pressure.^34,35^ To obtain beta stiffness, stress was initially calculated as *a=PD*/2WT, where P is the pressure in dynes/cm^2^, *D* is the lumen diameter, and WT is the wall thickness. Strain was further calculated as *E= {D-D*_*i*_*)/D*_*i*_, where *D*_*i*_ is the initial starting diameter. Data for each artery were fit to the curve using *a= a*_*i*_*e*^*6E*^ and *6* is the slope of tangential elastic modulus versus stress. A higher represents a stiffer artery.^36-38^

### Artery Histology

Sections of the thoracic aorta and middle cerebral arteries {MCAs} were frozen in optimal cutting temperature {OCT} compound and sliced into 8 µm sections using a cryostat {Leica, Wetzlar, Germany} and adhered to a charged slide. Both aortas and MCAs were blocked with 20% goat serum for 15 minutes in a humidified chamber, followed by an hour incubation with an anti-collagen-I, anti-collagen-III, or anti-a-smooth muscle actin primary antibody {ab270993 1:100, ab7778 1:50, ab124964 1:100}. The slides were further washed with PBS and incubated with a species-specific fluorescent secondary antibody {1:1000, 1:1000, 1:500}. Slides were washed and stained with DAPI {Life Tech, #P36935}. The slides were imaged with an Olympus Fluoview FV1000 {Tokyo, Japan} confocal microscope or a DeltaVision Ultra {Cytivia, Marlborough, MA} widefield microscope with deconvolution. Elastin was visualized by autofluorescence. Elastin, collagen I, collagen III, and a-smooth muscle actin content were analyzed by the mean gray intensity using FIJI by ImageJ software.

### Gene Expression

In samples of thoracic aortic arteries, mRNA gene expression was quantified for *COLlAl, COL3Al, COL4Al, COL4A2, MMP9, PDGFRA, and TGFBl* through qPCR. In samples of cerebral arteries {combined middle cerebral artery, posterior cerebral artery, and basilar artery} mRNA gene expression was quantified for *COLlAl, COL3Al, COL4Al, and COL4A2* through qPCR {see supplement Table for primer sequences}. RNA from thoracic aortic arteries and cerebral arteries were isolated by a standard protocol using Qiazol and RNAeasy Mini and Micro Kits {Qiagen, Hilden, Germany}. Isolated RNA was quantified using a NanoDrop 2000 {Thermo Scientific, Waltham, MA} with further reverse transcription performed to produce cDNA with a Qiagen QuantiTect Reverse Transcription Kit. The cDNA samples underwent real-time qPCR with ThermoFisher PowerUp Sybr Green using a QuantStudio 5 real-time PCR system {ThermoFisher Scientific, Waltham, MA}. mRNA expression was calculated using the 2^-CT^ method and 18s mRNA was used a housekeeping gene control with the values normalized to the young male group. Primers used are organized in Supplemental Table 2.

### Statistical Analysis

Statistical analyses were performed with IBM SPSS {Version 26, Armonk, NY} and GraphPad Prism 9.0. A two-way analysis of variance {ANOVA} between age and sex was used unless otherwise noted. In case of a significant F-value, post-hoc analyses were performed using a Tukey correction for preplanned comparisons. A repeated measures ANOVA was used to determine group differences for dose responses. Significance was set at p<0.05, and values are represented as mean ± SEM. Outliers were identified as z score>2 and removed from the data set.

## RESULTS

### Cerebral Artery Stiffness

In isolated PCAs incubated in a calcium-free solution, the changes in wall thickness and lumen diameter with increasing pressure were used to create stress-strain curves {Fig 1A, 1B}, and these curves were used to calculate -stiffness. For PCA passive -stiffness, there was an age effect {p= 0.02} with older PCAs having higher stiffness, but no sex differences {p=0.36, Fig 1C}. Similarly, aortic pulse wave velocity {PWV} was higher in old mice {p=0.04} but was similar between sexes {p=0.98, Fig 1D}.

**Figure 1.**
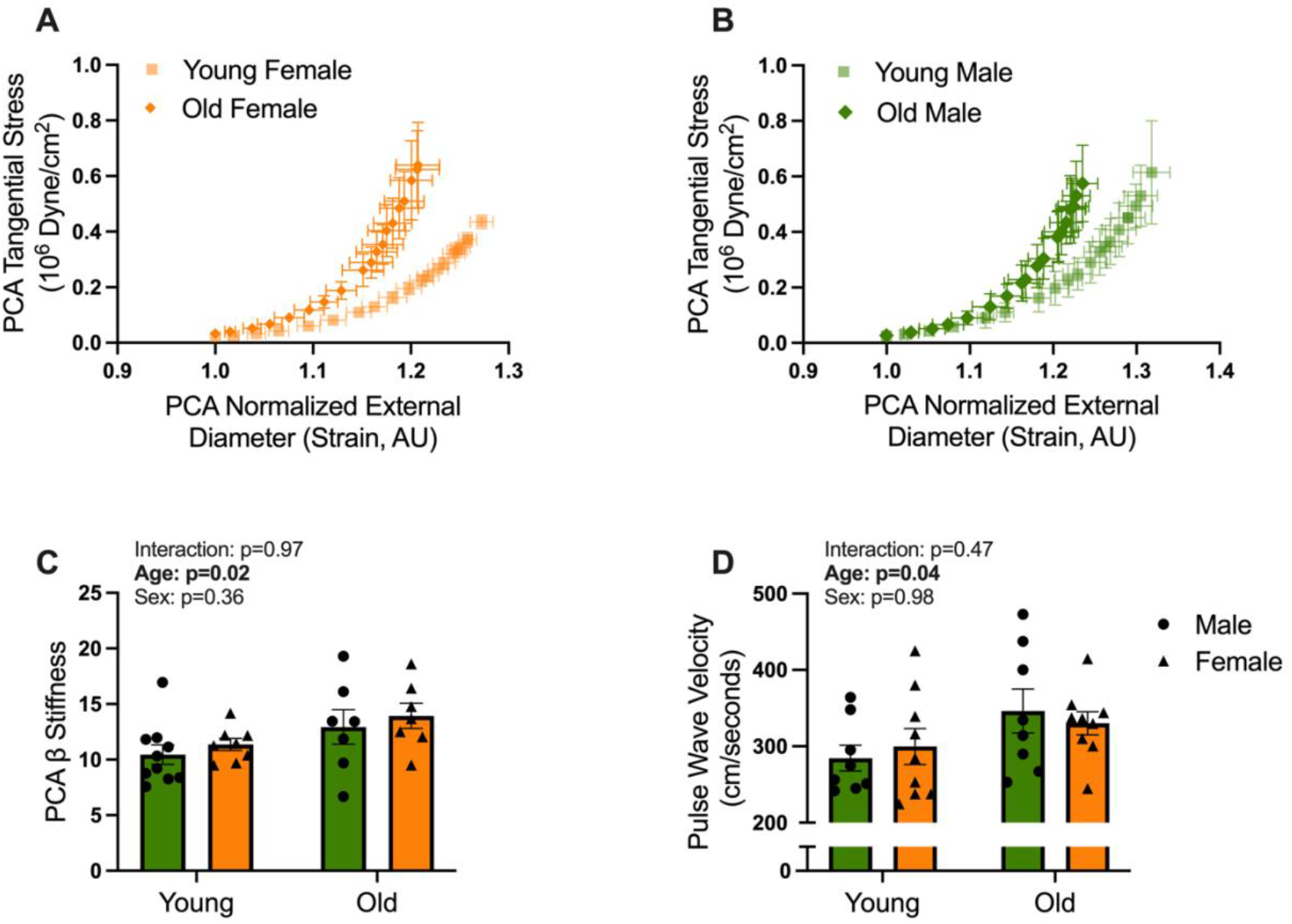
Cerebral artery passive stiffness and aortic stiffness are greater in old mice. E*x vivo* passive strain curves of posterior cerebral arteries {PCAs} are shown for *A)* young and old females and *B)* young and old males. *C) Ex vivo* PCA passive stiffness and *D)* aortic pulse wave velocity {PWV} measured in young and old females and males. *n*= 8-10/group. A two-way ANOVA with Tukey’s multiple comparisons was used. Data are mean ± SEM.

The active components of PCA stiffness were calculated from the changes in diameter during the pulsatile pressure application {with arteries bathed in a solution containing calcium}. During the low PP application, there was a main effect of sex on PCA compliance, stiffness, and distension {all p<0.05, Fig 2A, C, Supplemental Fig 1A, B}. In particular, old females had higher compliance and distension, as well as lower -stiffness compared with old males for the PCA during low PP application {all p<0.05}. During high PP application, there was also a main effect of sex on PCA compliance and distension {p=0.001, p=0.002 Fig 2E, F}. Unlike at low PP, the sex differences for compliance and distension at high PP were driven by the young groups, with higher distension in young females compared with young males {p= 0.004, Fig 2E; p=0.005, Fig 2F}. In addition, during the high PP condition, there was a main effect of age on PCA stiffness, compliance, and distension {all p<0.05, Fig 2D, E, F Supplemental Fig 1C, D}. Specifically, young females also had a higher PCA compliance and distension than the old females {all p< 0.01, Fig 2D, 2E, 2F}, but there were no age-related differences for compliance or distension in the male PCAs. These results suggest that the age-related increase in cerebral artery stiffness is more marked in females than males, at least during active conditions with high PP present.

**Figure 2.**
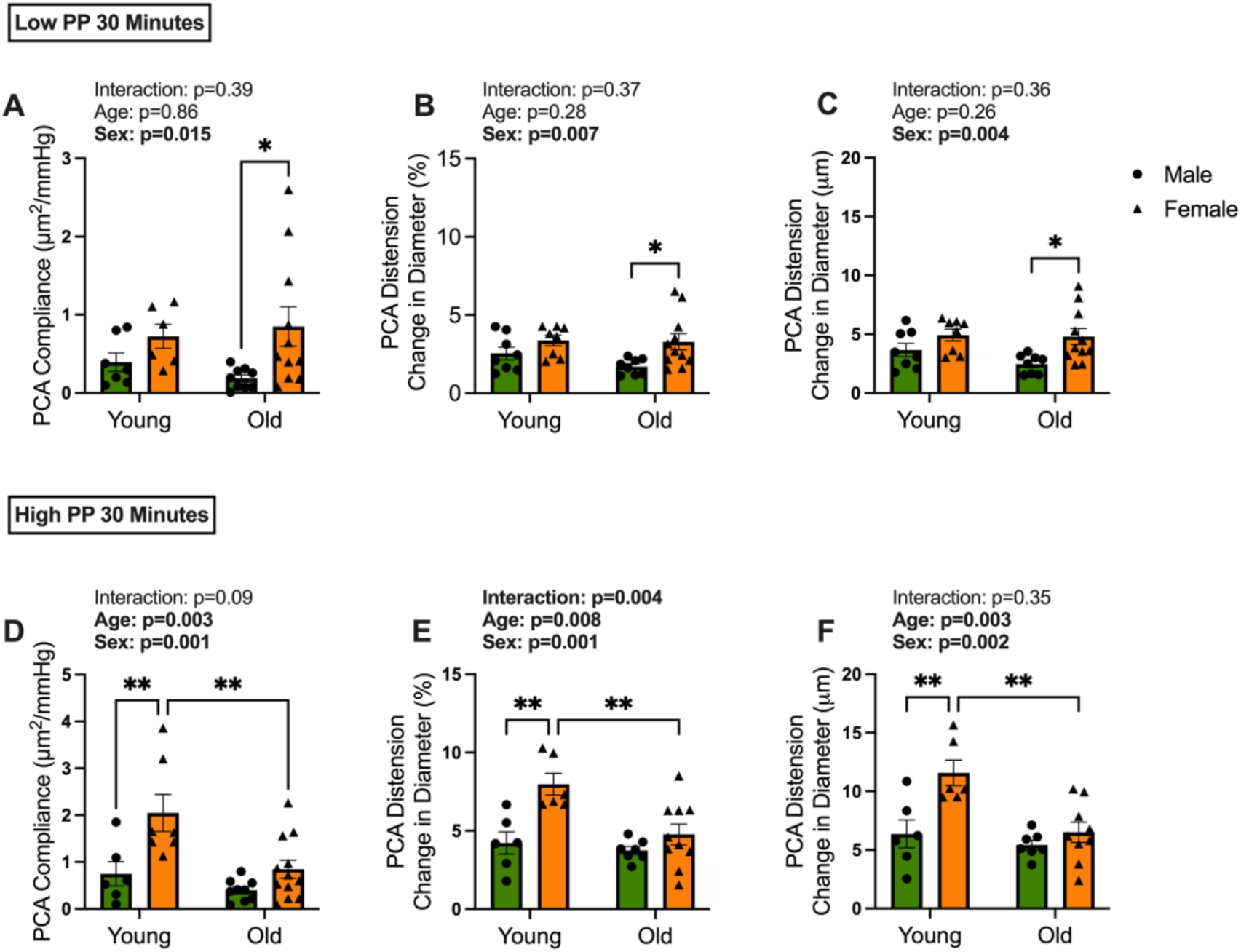
The decline in cerebral artery compliance with aging is greater in females. For young and old males and females, posterior cerebral artery {PCA} compliance during the application of A} low pulse {PP} and D} high PP, PCA percent distension {change in diameter} during B} low PP and E} high PP, and PCA absolute distention during C} low PP and F} high PP. n= 6-11/group. *\*P* < 0.05; *\*\*P* < 0.01. A two-way ANOVA with Tukey’s multiple comparisons was used. Data are mean ± SEM.

**Figure 3.**
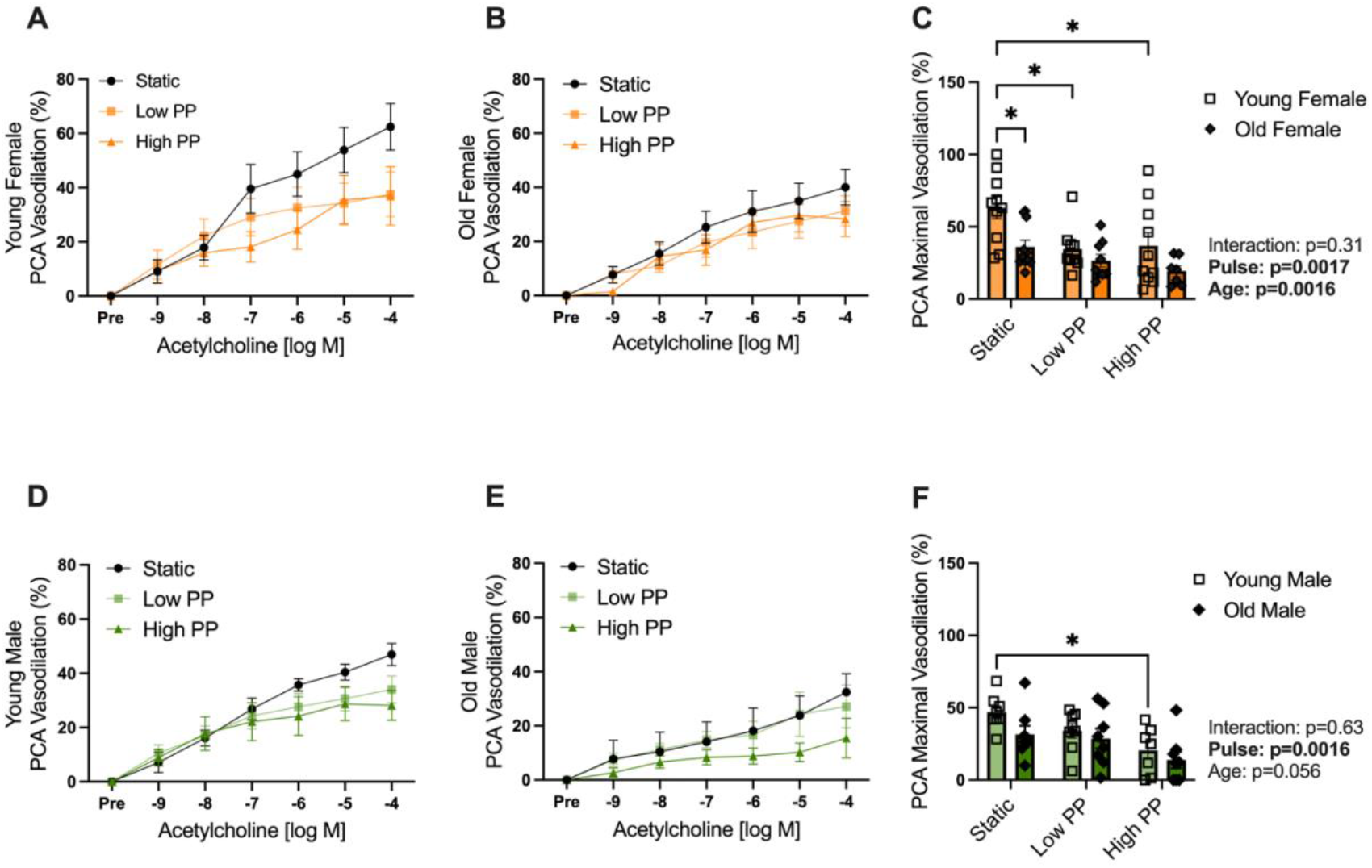
Low and high pulse pressure (PP) impairs endothelial function in cerebral arteries from young female mice. Endothelium-dependent dilation in the posterior cerebral arteries {PCAs} of *A)* young females, *B)* old females, *D)* young males, and *E)* old males to increasing doses of acetylcholine following exposure of static pressure, low PP, and high PP. Maximal vasodilation to acetylcholine for *C)* young and old female and *F)* young and old male PCAs following static pressure, low PP, and high PP. *n*= 6-11/group. *\*P <* 0.05. A two-way ANOVA with Tukey’s multiple comparisons was used. Data are mean ± SEM.

### Cerebral Artery Endothelial Function After Pulsatile Pressure Exposure

The effect of low and high PP was measured by endothelium-dependent vasodilation response to ACh in the PCAs. In PCAs from young female mice, the interaction effect of dose x pulse conditions was significant {p=0.002, Fig 1A}. The effect of the pulse condition was such that exposure to low and high PP reduced the maximal vasodilation to ACh when compared with the static PP condition {p= 0.02 and p= 0.04 respectively, Fig 1C}. In contrast to young females, the PCAs from old female mice had no interaction effect of dose x pulse conditions {p= 0.83, Fig 1B} or difference between PP conditions for the maximal PCA vasodilation to ACh {all p>0.05, Fig 1C}. During static pressure, PCAs from old females had lower maximal vasodilation to ACh compared with PCAs from young females {p=0.03, Fig 1C}. These findings demonstrate an impairment in ACh response with low and high PP in cerebral arteries from young females, but not old females.

An interaction of dose x pulse conditions {p=0.02, Fig 1D} was seen in young male PCAs. Specifically, the PCAs from young males had a lower maximal response to ACh after high PP when compared to static PP {p= 0.03, Fig 1F}. In PCAs from old male mice, there was no significant interaction of dose x pulse conditions {p= 0.57, Fig 1E} or difference between PP conditions for the maximal PCA vasodilation to Ach {all p>0.05, Fig 1F}. The findings demonstrate an impairment in ACh response with high PP in young male, but not old male cerebral arteries.

Endothelium-independent dilation, measured by dose-dependent vasodilation to sodium nitroprusside, was not different between young and old, female and male PCAs {interaction dose x group: p= 0.16, Supplemental Fig 2}. Thus, exposure to a low and PP resulted in impaired endothelium-dependent dilation in only young female cerebral arteries, while both young male and female cerebral arteries had impaired endothelium-dependent dilation after exposure to high PP.

### Aorta Wall Composition

The thickness of the aortic wall, both for the medial layer and the media+adventia, was significantly greater in old mice compared with young mice {p= 0.0005, p= 0.003, Fig 4A, B}. This difference with age was greatest in the males, with old males having a greater medial wall thickness in the aorta than young males {p= 0.01}, but the differences between the young and old females were not statistically significant {Fig 4B}. There were no differences in elastin content between all groups {Fig 4C}. There was an interaction of age x sex for aortic collagen I content, such that old males had lower collagen I content when compared to both young males {p= 0.0009, Fig 4D} and old females {p= 0.02, Fig 4D}. There were no age or sex effects for aorta collagen III content {Fig 4E}. There was lower a-smooth muscle actin with age in the aorta, which was particularly due to lower content in the old females when compared with the young females {p= 0.017, Fig 4F}. These results indicate that elastin remains unchanged, while, in contrast to previous studies^39^, aortic collagen I and a-smooth muscle actin decrease with aging in mice.

**Figure 4.**
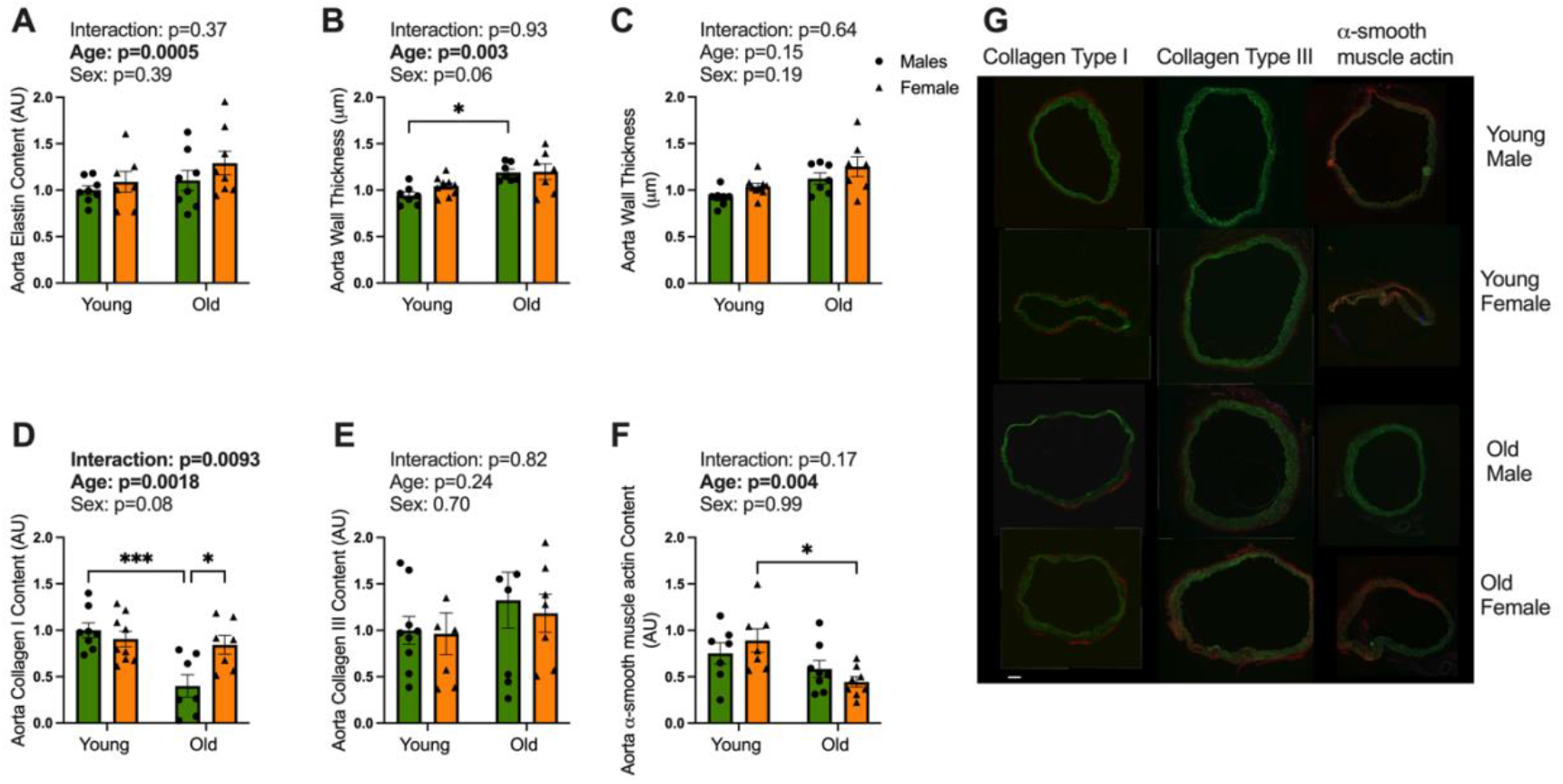
Old age is associated with greater wall thickness and less collagen and a-smooth muscle actin content in the aorta. Aortic artery *A)* wall thickness {medial layer} and *B)* wall thickness {medial and adventitial layer}, and content of *C)* elastin, *D)* collagen I, *E)* collagen III, and *F)* a-smooth actin measured by immunofluorescence in young and old females and males. Representative images are shown to the right: blue: DAPI, green: elastin, red: protein of interest. Scale bar: 100 µM *n*= 7-9/group. *\*P <* 0.05; *P <* 0.001. A two-way ANOVA with Tukey’s multiple comparisons was used. Data are mean ± SEM.

### Cerebral Artery Wall Composition

The impact of aging on the cerebral artery wall thickness was mediated by sex {interaction of age x sex, Fig 5A}. Specifically, old females had a greater cerebral artery wall thickness when compared with the young females {p= 0.016}, with no difference between the young and old males {Fig 5A}. In the cerebral arteries, there was an interaction of age x sex on elastin content, such that higher elastin content was observed for young males when compared with the young females {p= 0.0006} and old males {p= 0.0058, Fig 5B}. Cerebral artery collagen I content had a main effect of sex {p=0.0002} and was lower in males compared with females {p=0.04, Fig 5C}. There was no effect of sex or age on collagen III and a-smooth muscle actin in the cerebral arteries {Fig 5D, 5E}. These results indicate that age affects elastin content and wall thickness in the cerebral arteries but not the collagen or a-smooth muscle content. These results suggest that females have a higher cerebral artery collagen I content than males.

**Figure 5.**
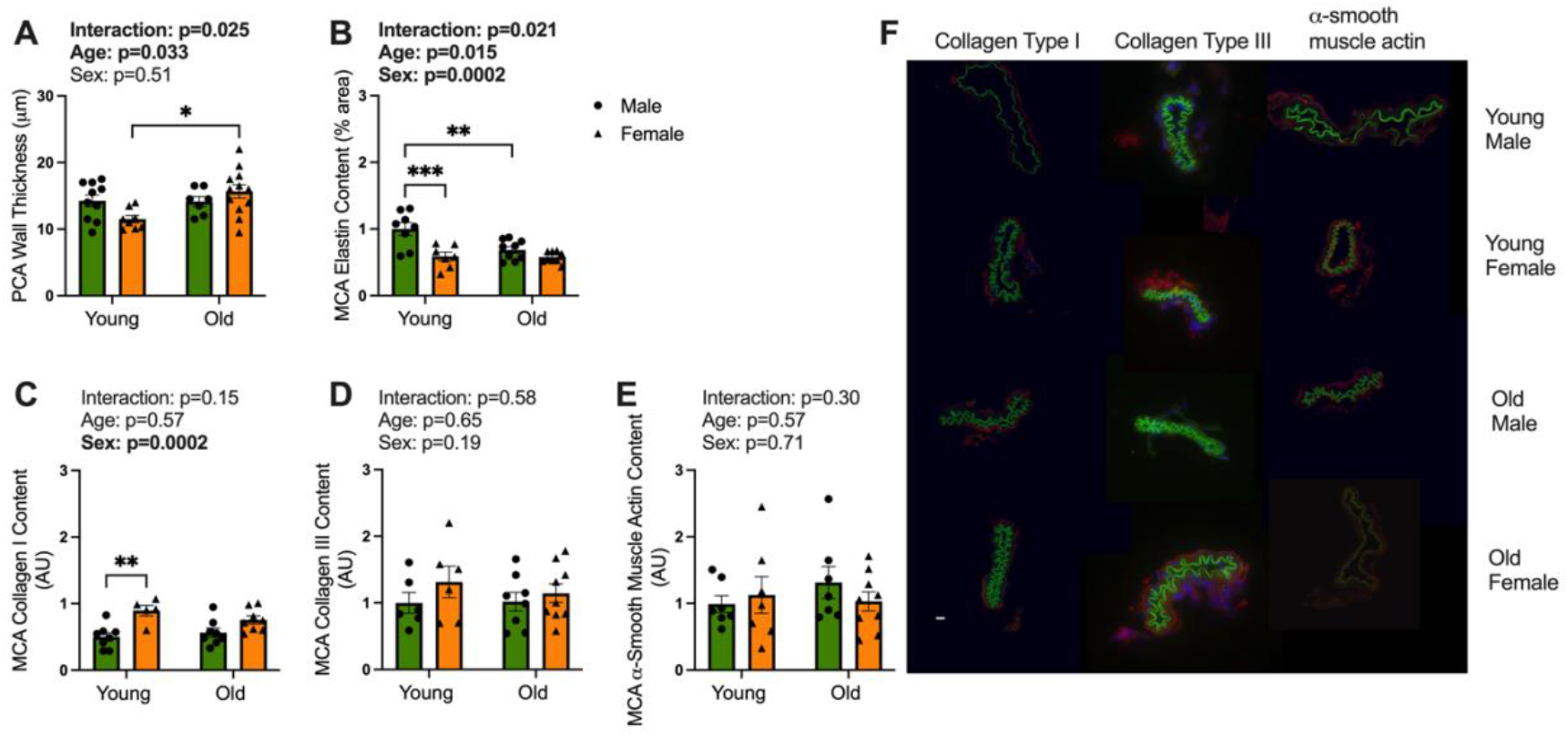
Age impacts cerebral artery wall thickness and elastin content, while sex impacts collagen content. *A)* Wall thickness of the posterior cerebral artery {PCA} measured by pressure myography at 50 mmHg. Content of *B)* elastin, *C)* collagen I, *D)* collagen III, and *E)* a-smooth actin measured by immunofluorescence in middle cerebral arteries {MCA} of young and old females and males. Representative images are shown to the right: blue: DAPI, green: elastin, red: protein of interest. Scale bar: 10 µM *n*= 6-10/group. *\*P <* 0.05; *\*\*P <* 0.01; *\*\*\*P <* 0.001. A two-way ANOVA with Tukey’s multiple comparisons was used. Data are mean ± SEM.

### Aorta Gene Expression

There was a main effect of age on *COLlAl, COL3Al, COL4Al, and COL4A2* gene expression in the aorta {all p<0.01}. Specifically, aorta *COLlAl, COL3Al*, and *COL4Al* gene expression were lower in old females when compared to young females {p= 0.01, p= 0.0007, p= 0.005}, while the expression did not differ between young and old males {Fig 6A, 6B, 6C}. Aorta gene expression of *COL4A2* was higher in young males compared to old males {p= 0.0003, Fig 6D}. Gene expression for *PDGFRa* and *TGFBl* did not differ between age or sex groups in the aortic artery {Fig 6E, 6F}. A main effect of age and sex was seen in aorta *MMP9* gene expression {all p<0.01}, with old females having higher expression compared to young females and old males {p= 0.05, p= 0.03, Fig 6G}.

**Figure 6.**
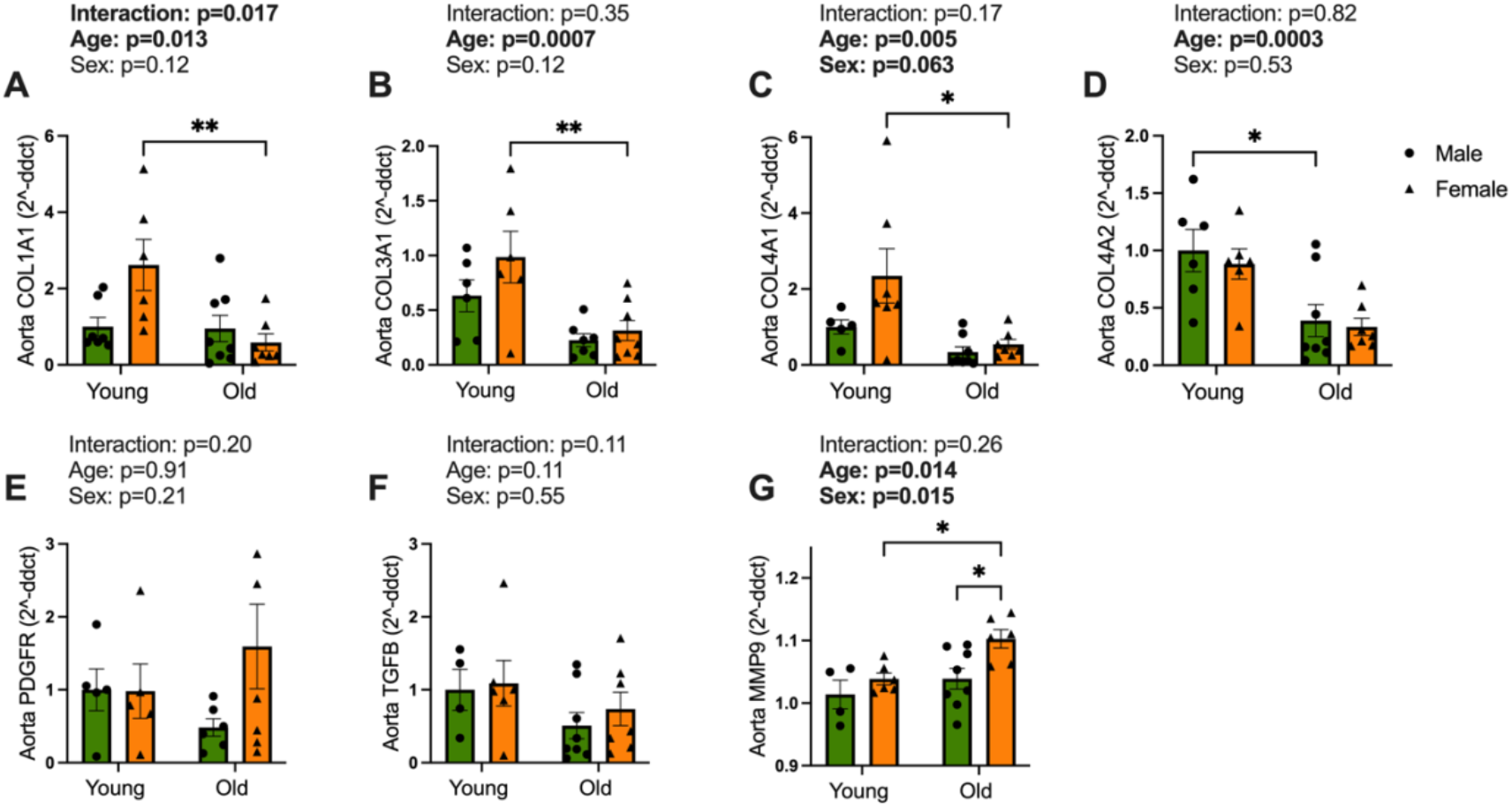
Collagen gene expression in the aorta is lower with old age. Aortic gene expression of *A) COLlAl* {collagen, type I, alpha 1 chain}, *B) COL3Al* {collagen, type III, alpha 1 chain}, *C) COL4Al* {collagen, type IV, alpha 1 chain}, *D) COL4A2* {collagen, type IV, alpha 2 chain}, *E) PDGFRa, F) TGFBl*, and *G) MMP9* in young and old females and males. *n*= 5-8/group. *\*P <* 0.05; *P <* 0.01. A two-way ANOVA with Tukey’s multiple comparisons was used. Data are mean ± SEM.

### Cerebral Artery Gene Expression

There was a difference in cerebral artery *COL3Al* gene expression with age {p=0.0012} and a trend for differences by sex {p=0.05. Fig 7B}. Specifically, young females had a greater cerebral artery *COL3Al* gene expression when compared with old females {p=0.05, Fig 7B}. Cerebral artery gene expression of *COL4Al* had a main effect of sex {p=0.03}, with the difference driven by young females having higher expression than young males {p=0.03, Fig 7C}. Cerebral artery gene expression for *COLlAl* and *COL4A2* did not differ with age or sex. {Fig 7A, D}.

**Figure 7.**
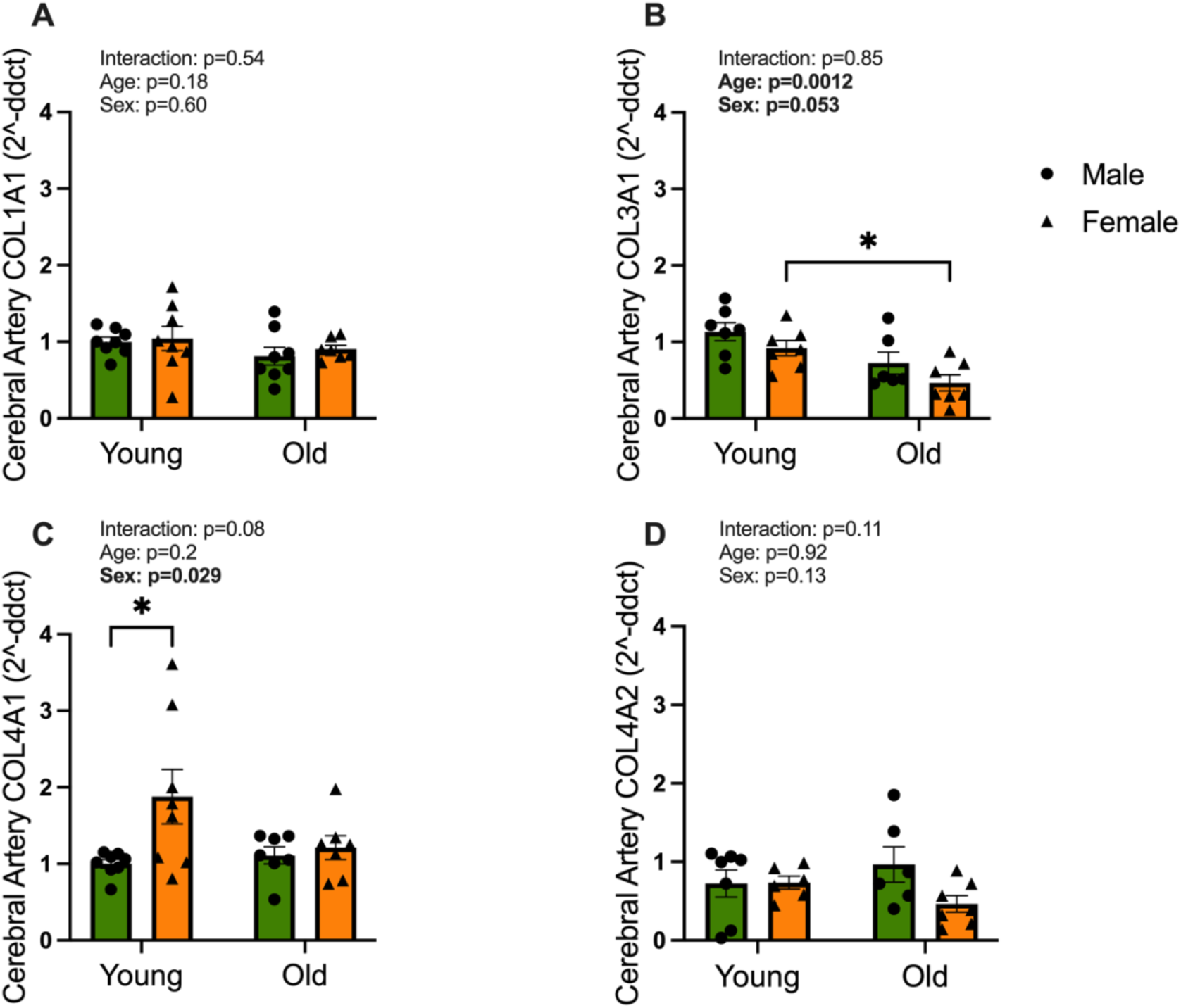
Cerebral artery gene expression for collagen is impacted by age and sex. Cerebral artery gene expression of *A) COLlAl* {collagen, type I, alpha 1 chain}, *B) COL3Al* {collagen, type III, alpha 1 chain}, *C) COL4Al* {collagen, type IV, alpha 1 chain}, *D) COL4A2* {collagen, type IV, alpha 2 chain}. *n*= 5-8/group. *\*P <* 0.05; *P <* 0.01. A two-way ANOVA with Tukey’s multiple comparisons was used. Data are mean ± SEM.

## DISCUSSION

The goal of this study was to determine the impact of age and sex on large and cerebral artery stiffness, as well as the cerebral artery susceptibility to damage by high PP. This study delivers novel evidence that young female mice have more compliant cerebral arteries and are thus more susceptible to high PP. The data indicate males and females undergo distinct extracellular matrix remodeling with aging, which is associated with changes in matrix metalloproteinases. Understanding how sex impacts age-related changes to vascular structure and function is an essential first step that has implications for our understanding of sex differences in disease risk.

### Impaired Cerebral Artery Endothelial Function with Advancing Age

Cerebrovascular function declines with age due, at least in part, to impairments in endothelial cell function.^40-42^ In this study we found impaired cerebral artery endothelial function with advancing age in female mice, similar to previous studies. ^43,44^ The decline in acetylcholine-mediated vasodilation with age was due to endothelial dysfunction as we found no differences in endothelium-independent dilation between age groups. Previous studies demonstrated that the age-related declines in cerebral endothelial function are due to greater oxidative stress and lower nitric oxide bioavailability.^20,41,45^ In this study, the age-related impairment in cerebral artery endothelium-dependent vasodilation did not reach statistical significance in the males, unlike our previous studies demonstrating differences between young and old male mice. ^20,43,46^ This discrepancy is partly due to the number of statistical comparisons made in this study, which reduced our power to detect differences. In addition, the ACh response was lower for the young males in this study than in our previous studies. In the previous studies, our mice were fed a diet containing a high amount of soybean meal, whereas in the current study, the mice were fed a soy-free diet. We have previously shown the protective effects of dietary soy on cerebral artery function.^47^ Our findings that young females have higher cerebral artery endothelial function than males align with the human research. Previous studies in humans find that young females have higher cerebral blood flow than young males, with the sex differences diminishing with age.^48^ Thus, the higher cerebral blood flow in young females may result from the greater cerebral artery endothelial function.

### High Pulse Pressure and Cerebral Artery Endothelial Dysfunction

As large arteries stiffen with advancing age, they cannot damp increased PP, and this increase in PP is associated with cerebrovascular impairments.^49^ In our study, compared with static pressure, cerebral arteries from young males and females had impaired endothelium-dependent vasodilation after exposure to high PP. This is similar to previous studies by us and others in young male arteries, where high PP impairs the acetylcholine response.^20,50^ It should be noted that this effect may be specific to acetylcholine-mediated vasodilation because PP improved the response to shear stress-induced vasodilation in young male cerebral arteries.^50^ However, our study is the first to examine the effects of pulsatile pressure in female cerebral arteries. While low PP does not impair young male cerebral arteries, we find that young female cerebral arteries have impaired endothelial function after low PP exposure. This impairment with low PP could be related to the higher compliance in the cerebral arteries from young female mice, leading to more wall stretch with PP exposure. This may suggest that female cerebral arteries are more susceptible to the increases in PP that occur with aging.

We also found that the cerebral artery response to PP was age dependent. Unlike young arteries, high PP did not yield lower endothelium-dependent dilation in old male or female cerebral arteries. While the existing impaired endothelium-dependent dilation at static pressure may play a role, we previously showed that the high PP response was significantly reduced with L-NAME in old animals after high PP exposure^20^, suggesting this was not a floor effect. Rather, the lack of an effect of high PP may be related to the higher stiffness and less distension of the old cerebral arteries during pulsatile pressure exposure. Overall, these results indicate that young cerebral arteries are susceptible to the negative effects of high PP, while old cerebral arteries are not impacted by high PP.

### The Association of Sex and Age with Cerebral Artery Stiffness

Increased cerebral artery stiffness with age could have devastating consequences on the brain. With stiffer cerebral arteries, elevated PP is delivered further into the cerebrovascular tree, potentially damaging the microvasculature.^1^ Greater stiffness in the cerebral arteries could also lead to decreased cerebral blood flow and impaired glymphatic clearance.^51,52^ In this study, the lack of an effect on high PP on cerebral arteries from old male and female mice suggests potential age-related changes to the arteries that are adaptive. A possible modification may be the increased stiffness seen in aged animals that protects the artery wall against added stretch. At young ages, we found no difference in passive stiffness between male and female cerebral arteries, similar to previous studies in arterioles.^53^ Furthermore, old cerebral arteries had a higher passive stiffness than young, and the age-related stiffening was similar between male and female mice. Previous studies have also reported greater passive stiffness in old cerebral arterioles compared with young.^30^ Concomitant with the higher stiffness in the cerebral arteries of old mice, we also found lower amounts of elastin, at least in males. However, unlike a previous study in arterioles,^30^ we did not find an age-related difference in collagen in the cerebral arteries, perhaps due to differences in measurement techniques. Overall, our results indicate that cerebral arteries from old mice have greater passive stiffness.

By applying pulsatile pressure ex vivo, we were able to measure cerebral artery compliance in an active state {i.e., with the contribution of contractile properties to stiffness}. We find that females have greater cerebral artery compliance than males. In addition, we find that old females have lower cerebral artery compliance compared with young females, but there is no difference in compliance between young and old males. This is especially important as females have a faster rate of increase in large artery stiffness with menopause compared with the rate in age-matched males.^18,54^ In addition, females have lower cerebral artery pulsatility early in life, and this protection does not persist with age.^55^ Previous human studies have shown older females have a greater rate of decline in cerebral artery blood velocity and a greater rate of increase of flow pulsatility when compared to older males.^55^ Thus, the greater cerebral artery compliance in young females may protect the microvasculature and other brain tissue, while also creating a situation where the arteries are more vulnerable to age-related increases in PP.

### Structural Differences of the Aorta with Sex and Age

When compared with cerebral arteries, much more is known about the structural changes to the aorta with aging. Aligned with previous studies, we find that aortic stiffness, measured by PWV, is greater in old age for both males and females.^46,56,57^ The age-related increase in aortic stiffness is likely due to changes to the extracellular matrix. The bulk of the collagen in the extracellular matrix is collagen I and III, with collagen I being responsible for providing mechanical strength and contractility^58^, while collagen III contributes to tensile strength and extensibility of the vessel wall.^59,60^ As a basement membrane protein, collagen IV is highly abundant in the artery walls and deficiency promotes vascular diseases, such as abdominal aortic aneurysm, dementia, and small vessel disease.^61,62^ The collagen content of the aortic wall during aging has been the subject of conflicting results. While some results show decreased arterial collagen content, others have reported increased collagen content with age.^39,63-65^ In our study, we found that aortic collagen I content was lower in old males compared with young males, but no differences in collagen III content. The distribution of collagen in the aorta is a major consideration as well; collagen I is prevalent in the adventitia, whereas collagen III comprises the majority of the medial layer.^64^ We performed our analysis of the entire aorta, not by region, which could explain the disparity in our results with previous studies.^39,63^

We also find a contrast between the collagen protein and gene expression, which may be because gene expression reflects the synthetic rate. Since collagen is a long-lived protein^66^, the synthetic rate does not necessarily align with the protein content. Older animals may produce less collagen and have a low turnover rate compared to younger animals.^64^ The higher expression of *MMP9* genes in older animals could explain the lower collagen content in the aorta. In addition, the formation of advanced-glycation-endproducts {AGEs} is increased with age.^67,68^ AGEs can form crosslinks connecting the polypeptide chains of the collagen fibrils, contributing to artery stiffness.^69,70^ Thus, even without an increase in collagen content, the large arteries in old animals may be stiffer due to the accumulation of collagen cross-links. In summary, we find that old mice have a stiffer aorta, but unlike previous studies, we do not find a concomitant increase in collagen content with age.

### Extracellular Matrix Difference of the Cerebral Vessels with Sex and Age

Along with the structural change in the aorta, our results indicate sex and age affect the structure of the cerebral vasculature. Mirroring the aorta, the collagen I content in old male cerebral arteries was lower compared to the young males, and *COL3Al* expression was lower in old female cerebral arteries compared to the young females. Interestingly, we also found that young female cerebral arteries had a higher expression of *COL4Al* than in young males, but old females and males were similar. Abnormalities in collagen IV have been linked to dementia and lower collagen IV is associated with lower cognitive function in animal models.^14^ As collagen IV plays a key role in the basement membrane and blood-brain-barrier formation, insufficient amounts of collagen IV can cause narrowing of the cerebral arteries to produce lower CBF.^71^ Previous studies found that mice with induced carotid stiffness had a diminished number of cerebral arteries containing collagen IV.^14^ The higher expression of *COL4Al* in young females may be what protects from cerebrovascular diseases.

## Limitations

There are several limitations to this study, the first being the lack of blood pressure measurements *in vivo*. The young females in our study showed an impairment in endothelium-dependent dilation to both low and high PP. Young females may have a lower starting PP *in vivo* compared to young males, thus explaining the vascular impairment seen with low PP. Second, the collagen content in both large and cerebral arteries was analyzed by immunofluorescent staining. Since this method is only semi-quantitative, our disparate findings from previous studies should be taken cautiously.^39^ Third, we did not account for the estrous cycle in female mice. However, in our previous study, we demonstrated that although the estrous cycle affects *in vivo* arterial stiffness, artery endothelial function in isolated cerebral arteries was unaffected.^72^ In addition, female mice do not undergo a menopause that is similar to humans, instead female mice experience either an estrous pause, where the mouse experiences irregular estrous cycles, or transition into an anestrous state, where ovulatory cycles halt with low levels of sex steroids present. ^73,74^ Therefore, our findings in old female mice may be less translatable to humans than the findings in old males.

## Conclusion

Our findings add to our understanding of age and sex-related differences in cerebral artery structure and the consequences of PP on endothelial function. As old age leads to increased stiffness, the vasculature may readily adapt to the exposure of PP, as shown by the lack of dysfunction in endothelium-dependent dilation after exposure to PP for both old male and female mice. However, the negative consequences of increased stiffness are ever present, including potential damage to the organs with high blood flow, such as the brain. We also found that young female mice had more compliant cerebral arteries that were more susceptive to dysfunction caused by both low and high PP. Consequently, females are at a higher risk for a decline in cerebrovascular function during the increases in PP with aging. This may contribute to the higher risk of neurodegenerative and cerebrovascular diseases in females after menopause.

## Sources of Funding

This work was supported by National Institutes of Health R01 AG064016.

## Authors’ Contributions

YC and AW contributed to study concept and design. YC, JS, PK, AC, JL, and AW contributed to data collection and analysis. YC and AW contributed to development of figures/tables. YC and AW drafted the manuscript, and all other authors edited and approved final manuscript.

## Declaration of Conflicting Interests

The author{s} declared no potential conflicts of interest with respect to the research, authorship, and/or publication of this article.

## Notes

### Competing Interest Statement

The authors have declared no competing interest.

## CITATIONS

1. Mitchell, G. F. Aortic stiffness, pressure and flow pulsatility, and target organ damage. J. Appl. Physiol. Bethesda Md 1985 125, 1871–1880 (2018).

2. Pase, M. P., Herbert, A., Grima, N. A., Pipingas, A. & O’Rourke, M. F. Arterial stiffness as a cause of cognitive decline and dementia: a systematic review and meta-analysis. Intern. Med. J. 42, 808–815 (2012).

3. Hughes, T. M. et al. Arterial stiffness and dementia pathology. Neurology 90, e1248–e1256 (2018).

4. Peng, S.-L. et al. Age-related changes in cerebrovascular reactivity and their relationship to cognition: A four-year longitudinal study. NeuroImage 174, 257–262 (2018).

5. Qiu, C., Winblad, B., Viitanen, M. & Fratiglioni, L. Pulse Pressure and Risk of Alzheimer Disease in Persons Aged 75 Years and Older. Stroke 34, 594–599 (2003).

6. Mitchell, G. F. Effects of central arterial aging on the structure and function of the peripheral vasculature: implications for end-organ damage. J. Appl. Physiol. 105, 1652–1660 (2008).

7. Tarumi, T., Shah, F., Tanaka, H. & Haley, A. P. Association Between Central Elastic Artery Stiffness and Cerebral Perfusion in Deep Subcortical Gray and White Matter. Am. J. Hypertens. 24, 1108–1113 (2011).

8. Rensma, S. P. et al. Associations of Arterial Stiffness With Cognitive Performance, and the Role of Microvascular Dysfunction. Hypertension 75, 1607–1614 (2020).

9. Jefferson, A. L. et al. Higher Aortic Stiffness Is Related to Lower Cerebral Blood Flow and Preserved Cerebrovascular Reactivity in Older Adults. Circulation 138, 1951–1962 (2018).

10. Rosano, C. et al. Aortic Pulse Wave Velocity Predicts Focal White Matter Hyperintensities in a Biracial Cohort of Older Adults. Hypertension 61, 160–165 (2013).

11. Pase, M. P. et al. Association of Aortic Stiffness With Cognition and Brain Aging in Young and Middle-Aged Adults: The Framingham Third Generation Cohort Study. Hypertens. Dallas Tex 1979 67, 513–519 (2016).

12. Walker, A. E. et al. Greater impairments in cerebral artery compared with skeletal muscle feed artery endothelial function in a mouse model of increased large artery stiffness. J. Physiol. 593, 1931–1943 (2015).

13. Sadekova, N., Vallerand, D., Guevara, E., Lesage, F. & Girouard, H. Carotid Calcification in Mice: A New Model to Study the Effects of Arterial Stiffness on the Brain. J. Am. Heart Assoc. Cardiovasc. Cerebrovasc. Dis. 2, e000224 (2013).

14. Muhire, G. et al. Arterial Stiffness Due to Carotid Calcification Disrupts Cerebral Blood Flow Regulation and Leads to Cognitive Deficits. J. Am. Heart Assoc. Cardiovasc. Cerebrovasc. Dis. 8, e011630 (2019).

15. Beam, C. R. et al. Differences Between Women and Men in Incidence Rates of Dementia and Alzheimer’s Disease. J. Alzheimers Dis. JAD 64, 1077–1083 (2018).

16. Lin, K. A. et al. Marked gender differences in progression of mild cognitive impairment over 8 years. Alzheimers Dement. N. Y. N 1, 103–110 (2015).

17. Coutinho, T. Arterial Stiffness and Its Clinical Implications in Women. Can. J. Cardiol. 30, 756–764 (2014).

18. DuPont, J. J., Kim, S. K., Kenney, R. M. & Jaffe, I. Z. Sex differences in the time course and mechanisms of vascular and cardiac aging in mice: role of the smooth muscle cell mineralocorticoid receptor. Am. J. Physiol.-Heart Circ. Physiol. 320, H169–H180 (2021).

19. Hawes, J. Z. et al. Elastin haploinsufficiency in mice has divergent effects on arterial remodeling with aging depending on sex. Am. J. Physiol. - Heart Circ. Physiol. 319, H1398–H1408 (2020).

20. Winder, N. R. et al. High pulse pressure impairs cerebral artery endothelial function in young, but not old, mice. Exp. Gerontol. 173, 112101 (2023).

21. Dingemans, K. P., Teeling, P., Lagendijk, J. H. & Becker, A. E. Extracellular matrix of the human aortic media: An ultrastructural histochemical and immunohistochemical study of the adult aortic media. Anat. Rec. 258, 1–14 (2000).

22. Tsamis, A., Krawiec, J. T. & Vorp, D. A. Elastin and collagen fibre microstructure of the human aorta in ageing and disease: a review. J. R. Soc. Interjace 10, 20121004 (2013).

23. Lakatta, E. G. & Levy, D. Arterial and cardiac aging: major shareholders in cardiovascular disease enterprises: Part I: aging arteries: a ‘set up’ for vascular disease. Circulation 107, 139–146 (2003).

24. de Montgolfier, O. et al. High Systolic Blood Pressure Induces Cerebral Microvascular Endothelial Dysfunction, Neurovascular Unit Damage, and Cognitive Decline in Mice. Hypertension 73, 217–228 (2019).

25. Zhu, W. et al. TGF1 reinforces arterial aging in the vascular smooth muscle cell through a long-range regulation of the cytoskeletal stiffness. Sci. Rep. 8, 2668 (2018).

26. Hu, Y., Bock, G., Wick, G. & Xu, Q. Activation of PDGF receptor alpha in vascular smooth muscle cells by mechanical stress. FASEB J. Off. Publ. Fed. Am. Soc. Exp. Biol. 12, 1135–1142 (1998).

27. Podzolkov, V. I., Nebieridze, N. N. & Safronova, T. A. Transforming Growth Factor-1, Arterial Stiffness and Vascular Age in Patients With Uncontrolled Arterial Hypertension. Heart Lung Circ. 30, 1769–1777 (2021).

28. Brown, X. Q. et al. Effect of substrate stiffness and PDGF on the behavior of vascular smooth muscle cells: implications for atherosclerosis. J. Cell. Physiol. 225, 115–122 (2010).

29. Xu, X. et al. Recent Progress in Vascular Aging: Mechanisms and Its Role in Age-related Diseases. Aging Dis. 8, 486–505 (2017).

30. Diaz-Otero, J. M., Garver, H., Fink, G. D., Jackson, W. F. & Dorrance, A. M. Aging is associated with changes to the biomechanical properties of the posterior cerebral artery and parenchymal arterioles. Am. J. Physiol.-Heart Circ. Physiol. 310, H365–H375 (2016).

31. Sabharwal, R., Chapleau, M. W., Gerhold, T. D., Baumbach, G. L. & Faraci, F. M. Plasticity of cerebral microvascular structure and mechanics during hypertension and following recovery of arterial pressure. Am. J. Physiol.-Heart Circ. Physiol. 323, H1108–H1117 (2022).

32. Flamant, M. et al. Role of matrix metalloproteinases in early hypertensive vascular remodeling. Hypertens. Dallas Tex 1979 50, 212–218 (2007).

33. Leloup, A. J. A. et al. A novel set-up for the ex vivo analysis of mechanical properties of mouse aortic segments stretched at physiological pressure and frequency. J. Physiol. 594, 6105–6115 (2016).

34. Iadecola, C. et al. Impact of Hypertension on Cognitive Function: A Scientific Statement From the American Heart Association. Hypertens. Dallas Tex 1979 68, e67–e94 (2016).

35. Modrick, M. L., Didion, S. P., Sigmund, C. D. & Faraci, F. M. Role of oxidative stress and AT1 receptors in cerebral vascular dysfunction with aging. Am. J. Physiol. Heart Circ. Physiol. 296, H1914–1919 (2009).

36. Nakano, T., Tominaga, R., Nagano, I., Okabe, H. & Yasui, H. Pulsatile flow enhances endothelium-derived nitric oxide release in the peripheral vasculature. Am. J. Physiol. Heart Circ. Physiol. 278, H1098–1104 (2000).

37. Noris, M. et al. Nitric oxide synthesis by cultured endothelial cells is modulated by flow conditions. Circ. Res. 76, 536–543 (1995).

38. Osei-Owusu, P. et al. Altered reactivity of resistance vasculature contributes to hypertension in elastin insufficiency. Am. J. Physiol. Heart Circ. Physiol. 306, H654–666 (2014).

39. Longtine, A. G. et al. Female C57BL/6N mice are a viable model of aortic aging in women. Am. J. Physiol.-Heart Circ. Physiol. 324, H893–H904 (2023).

40. Seals, D.R., Jablonski, K.L. & Donato, A.J. Aging and vascular endothelial function in humans. Clin. Sci. Lond. Engl. 1979 120, 357–375 (2011).

41. Walker, A. E. et al. Beneficial effects of lifelong caloric restriction on endothelial function are greater in conduit arteries compared to cerebral resistance arteries. Age Dordr. Neth. 36, 559–569 (2014).

42. Donato, A. J. et al. Direct evidence of endothelial oxidative stress with aging in humans: relation to impaired endothelium-dependent dilation and upregulation of nuclear factor-kappaB. Circ. Res. 100, 1659–1666 (2007).

43. Cole, J. A. et al. Sex Differences in the Relation Between Frailty and Endothelial Dysfunction in Old Mice. J. Gerontol. A. Biol. Sci. Med. Sci. 77, 416–423 (2022).

44. Takenouchi, Y., Kobayashi, T., Matsumoto, T. & Kamata, K. Gender differences in age-related endothelial function in the murine aorta. Atherosclerosis 206, 397–404 (2009).

45. Csiszar, A. et al. Caloric restriction confers persistent anti-oxidative, pro-angiogenic, and anti-inflammatory effects and promotes anti-aging miRNA expression profile in cerebromicrovascular endothelial cells of aged rats. Am. J. Physiol.-Heart Circ. Physiol. 307, H292–H306 (2014).

46. Reeve, E. H. et al. Pyridoxamine treatment ameliorates large artery stiffening and cerebral artery endothelial dysfunction in old mice. J. Cereb. Blood Flow Metab. 43, 281–295 (2023).

47. Kehmeier, M. N. et al. Effects of dietary soy content on cerebral artery function and behavior in ovariectomized female mice. Am. J. Physiol. Heart Circ. Physiol. 326, H636–H647 (2024).

48. Mazzucco, S., Li, L., Tuna, M. A. & Rothwell, P. M. Age-specific sex-differences in cerebral blood flow velocity in relation to haemoglobin levels. Eur. Stroke J. 9, 772–780 (2024).

49. Thorin-Trescases, N. et al. Impact of pulse pressure on cerebrovascular events leading to age-related cognitive decline. Am. J. Physiol. Heart Circ. Physiol. 314, H1214–H1224 (2018).

50. Raignault, A., Bolduc, V., Lesage, F. & Thorin, E. Pulse pressure-dependent cerebrovascular eNOS regulation in mice. J. Cereb. Blood Flow Metab. Off. J. Int. Soc. Cereb. Blood Flow Metab. 37, 413–424 (2017).

51. van den Kerkhof, M. et al. Impaired damping of cerebral blood flow velocity pulsatility is associated with the number of perivascular spaces as measured with 7T MRI. J. Cereb. Blood Flow Metab. Off. J. Int. Soc. Cereb. Blood Flow Metab. 43, 937–946 (2023).

52. Bown, C. W. et al. Enlarged perivascular space burden associations with arterial stiffness and cognition. Neurobiol. Aging 124, 85–97 (2023).

53. Chambers, L. C., Yen, M., Jackson, W. F. & Dorrance, A. M. Female mice are protected from impaired parenchymal arteriolar TRPV4 function and impaired cognition in hypertension. Am. J. Physiol.-Heart Circ. Physiol. 324, H581–H597 (2023).

54. Seeland, U., Demuth, I., Regitz-Zagrosek, V., Steinhagen-Thiessen, E. & Konig, M. Sex differences in arterial wave reflection and the role of exogenous and endogenous sex hormones: results of the Berlin Aging Study II. J. Hypertens. 38, 1040 (2020).

55. Alwatban, M. R. et al. Effects of age and sex on middle cerebral artery blood velocity and flow pulsatility index across the adult lifespan. J. Appl. Physiol. 130, 1675–1683 (2021).

56. Ahimastos, A. A., Formosa, M., Dart, A. M. & Kingwell, B. A. Gender Differences in Large Artery Stiffness Pre- and Post Puberty. J. Clin. Endocrinol. Metab. 88, 5375–5380 (2003).

57. Sutton-Tyrrell, K. et al. Elevated Aortic Pulse Wave Velocity, a Marker of Arterial Stiffness, Predicts Cardiovascular Events in Well-Functioning Older Adults. Circulation 111, 3384–3390 (2005).

58. Ponticos, M., Partridge, T., Black, C. M., Abraham, D. J. & Bou-Gharios, G. Regulation of Collagen Type I in Vascular Smooth Muscle Cells by Competition between Nkx2.5 and 6EF1/ZEB1. Mol. Cell. Biol. 24, 6151–6161 (2004).

59. de Souza, R.R. Aging of myocardial collagen. Biogerontology 3, 325–335 (2002).

60. Liu, X., Wu, H., Byrne, M., Krane, S. & Jaenisch, R. Type III collagen is crucial for collagen I fibrillogenesis and for normal cardiovascular development. Proc. Natl. Acad. Sci. 94, 1852–1856 (1997).

61. Lanfranconi, S. & Markus, H. S. COL4A1 mutations as a monogenic cause of cerebral small vessel disease: a systematic review. Stroke 41, e513–518 (2010).

62. Steffensen, L. B. et al. Basement membrane collagen IV deficiency promotes abdominal aortic aneurysm formation. Sci. Rep. 11, 12903 (2021).

63. Thompson, R. W. Reflections on the pathogenesis of abdominal aortic aneurysms. Cardiovasc. Surg. 10, 389–394 (2002).

64. Rahkonen, O. et al. Mice With a Deletion in the First Intron of the Col1a1 Gene Develop Age-Dependent Aortic Dissection and Rupture. Circ. Res. 94, 83–90 (2004).

65. Fleenor, B. S., Marshall, K. D., Durrant, J. R., Lesniewski, L. A. & Seals, D. R. Arterial stiffening with ageing is associated with transforming growth factor-1-related changes in adventitial collagen: reversal by aerobic exercise. J. Physiol. 588, 3971–3982 (2010).

66. Toyama, B. H. et al. Identification of long-lived proteins reveals exceptional stability of essential cellular structures. Cell 154, 971–982 (2013).

67. Monnier, V. M., Sell, D. R., Abdul-Karim, F. W. & Emancipator, S. N. Collagen browning and cross-linking are increased in chronic experimental hyperglycemia. Relevance to diabetes and aging. Diabetes 37, 867–872 (1988).

68. LaRocca, T. J., Martens, C. R. & Seals, D. R. Nutrition and other lifestyle influences on arterial aging. Ageing Res. Rev. 39, 106–119 (2017).

69. Fessel, G. et al. Advanced Glycation End-Products Reduce Collagen Molecular Sliding to Affect Collagen Fibril Damage Mechanisms but Not Stiffness. PLOS ONE 9, e110948 (2014).

70. Grandhee, S. K. & Monnier, V. M. Mechanism of formation of the Maillard protein cross-link pentosidine. Glucose, fructose, and ascorbate as pentosidine precursors. J. Biol. Chem. 266, 11649–11653 (1991).

71. Joutel, A. & Faraci, F. M. Cerebral small vessel disease: insights and opportunities from mouse models of collagen IV-related small vessel disease and cerebral autosomal dominant arteriopathy with subcortical infarcts and leukoencephalopathy. Stroke 45, 1215–1221 (2014).

72. Kehmeier, M. N. et al. In vivo arterial stiffness, but not isolated artery endothelial function, varies with the mouse estrous cycle. Am. J. Physiol. Heart Circ. Physiol. 323, H1057–H1067 (2022).

73. Koebele, S. V. & Bimonte-Nelson, H. A. Modeling menopause: The utility of rodents in translational behavioral endocrinology research. Maturitas 87, 5–17 (2016).

74. Huang, H.H., Steger, R.W., Bruni, J.F. & Meites, J. Patterns of Sex Steroid and Gonadotropin Secretion in Aging Female Rats*. Endocrinology 103, 1855–1859 (1978).

