## Supplemental Data for "The Impact of Age and Sex on Cerebral and Large Artery Stiffness and the Response to Pulse Pressure"

### Supplemental Table 1. Animal Characteristics

**Table 1: Animal Characteristics**

| Variable | Young Females | Old Females | Young Males | Old Males |
|---|---|---|---|---|
| N | 10 | 8 | 9 | 10 |
| Age, months | 6±1 | 24±0.8 | 6±0.8 | 24±0.6 |
| Body mass (g)[a,b,c] | 22.8 ± 1.29 | 32.3 ± 3.04* | 33.0 ±3.12* | 33.5±3.20* |
| Heart mass (mg) [a,b,c] | 0.12 ±0.02 | 0.19±0.04* | 0.16±0.02* | 0.18±0.01* |
| Percent heart:body mass[a] | 0.51±0.1 | 0.58±0.1 | 0.50±0.05† | 0.53±0.06 |
| Liver mass (mg)[b,c] | 1.06 ±0.38 | 1.69 ±0.51* | 1.95 ±0.48* | 1.69 ±0.62* |
| Percent liver:body mass | 4.63±1.59 | 5.26±1.63 | 5.88±1.16 | 5.04±1.82 |
| WAT mass (mg)[c] | 0.410±1.66 | 1.07±-0.54* | 1.02±0.50* | 0.770±0.46* |
| Percent WAT:body mass[c] | 1.78±0.64 | 3.03±1.51* | 3.03±1.33 | 2.21±1.12† |
| Spleen mass (mg)[a] | 0.74±0.025 | 0.18±0.10* | 0.09±0.026 | 0.12±0.12 |
| Percent spleen:body mass[a] | 0.33±0.11 | 0.56±0.33 | 0.28±0.066 | 0.352±0.39 |
| Gastroc mass (mg)[a] | 0.145±0.016 | 0.132±0.41 | 0.178±0.041 | 0.133±0.049*‡ |
| Percent gastroc:body mass[a] | 0.571±0.21 | 0.391±0.18* | 0.552±0.28 | 0.400±0.15‡ |
| Soleus mass (mg)[b,c] | 0.009±0.002 | 0.009±0.003 | 0.01±0.007*† | 0.01±0.002‡ |
| Percent soleus:body mass | 0.03±0.02 | 0.03±0.01 | 0.039±0.02† | 0.03±0.006‡ |
| PCA Maximum Diameter | 147.9±10.1 | 162.8±8.7* | 158.5±20.1 | 167.1±6.6* |
| Wall: Lumen [b] | 0.12±0.2 | 0.11±0.02 | 0.09±0.02* | 0.08±0.01*† |
| Uterus mass (mg)[a] | 0.09±0.02 | 0.1±0.06* | | |
| Percent uterus:body mass[a] | 0.4±0.1 | 0.2±0.2* | | |

Data are mean ± SEM. WAT, white adipose tissue. [a] p<0.05 main effect of age, [b] p<0.05 main effect of sex, [c] p<0.05 interaction age x sex, * P<0.05 vs. young females, † P<0.05 vs. old females, ‡ P<0.05 vs. young males

### Supplemental Table 2. Gene Expression Primer Sequence

| **Primer** | **Forward** | **Reverse** |
|---|---|---|
| *COL1A1* | CTGGCGGTTCAGGTCCAAT | TTCCAGGCAATCCACGAGC |
| *COL3A1* | CAGGACCTAAGGGCGAAGATG | TCCGGGCATACCCCGTATC |
| *COL4A1* | CCTGGCACAAAAGGGACGA | ACGTGGCCGAGAATTTCACC |
| *COL4A2* | GGACCCAAGGGACAACCAG | CCCAACAAGTGTGATGTCAGAT |
| *PDGFRA* | AGAGTTACACGTTTGAGCTGTC | GTCCCTCCACGGTACTCCT |
| *TGFB1* | CTCCCGTGGCTTCTAGTGC | GCCTTAGTTTGGACAGGATCTG |
| *MMP9* | GCAGAGGCATACTTGTACCG | TGATGTTATGATGGTCCCACTTG |
| *18s* | TAGAGGGACAAGTGGCGTTC | CGCTGAGCCAGTCAGTGT |

**Supplemental Figure 1.**

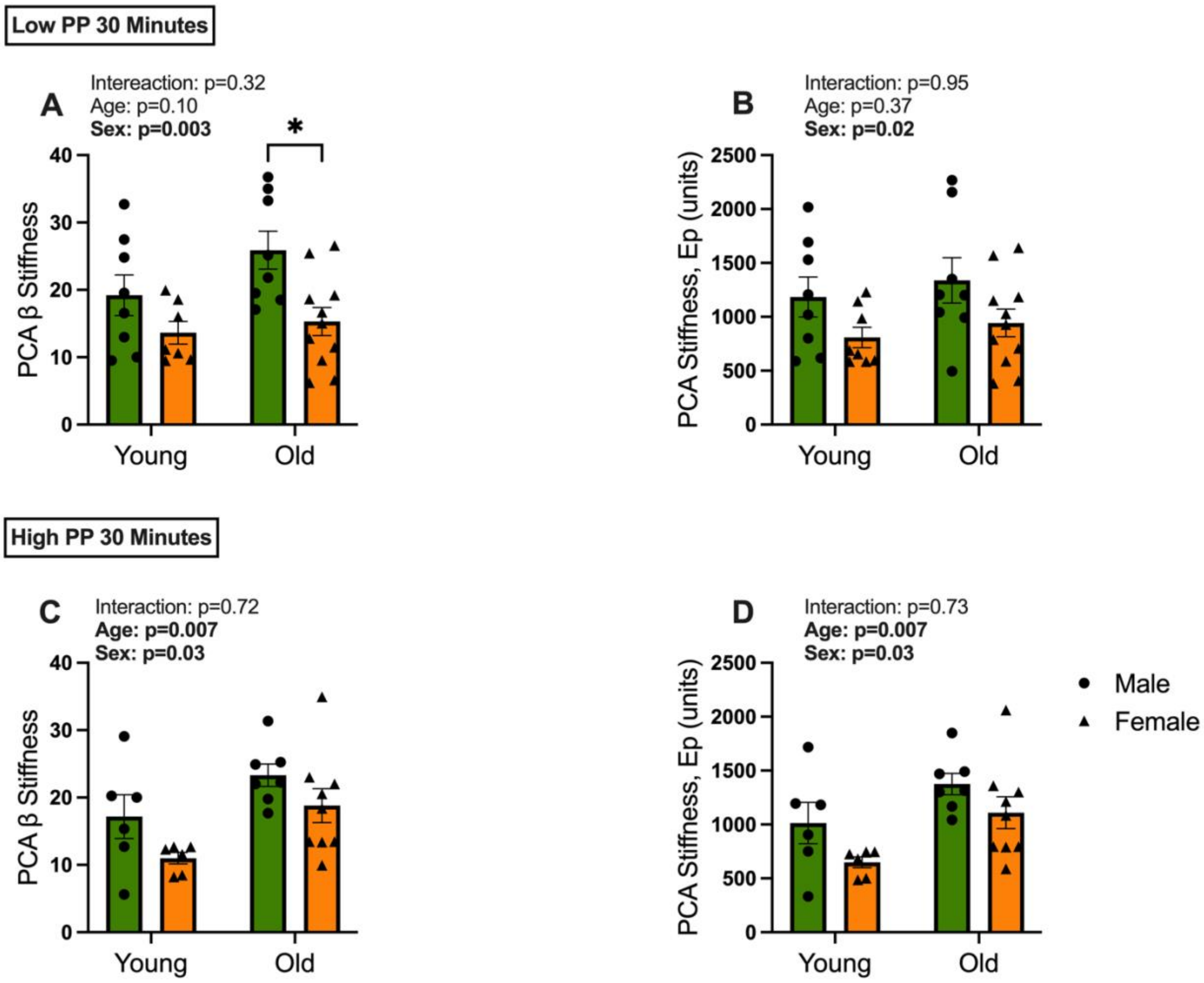


**Supplemental Figure 1. Cerebral artery stiffness is higher in both sexes with age post low and high PP.** PCA beta stiffness of posterior cerebral artery (PCA) from young and old, female and male mice during *A)* low pulse pressure (PP) and *B)* high pulse pressure (PP). PCA stiffness, EP of PCA for young and old, female and male mice during *E)* low PP and *F)* high PP. *n*= 6-11/group. **P* < 0.05; ***P* < 0.01. A two-way ANOVA with Tukey's multiple comparisons was used. Data are mean ± SEM.

**Supplemental Figure 2.**

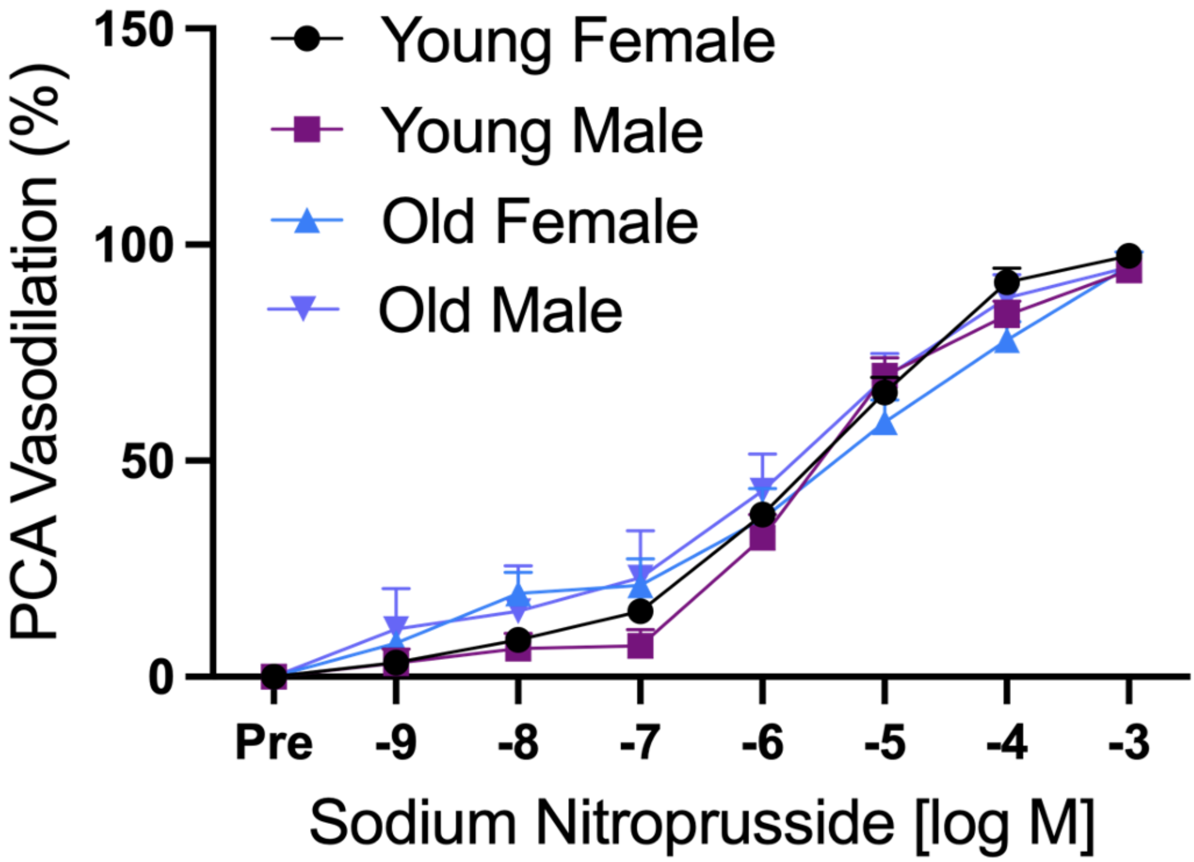


**Supplemental Figure 2.** Endothelium-independent dilation was maintained in the cerebral arteries of all groups. Dose dependent vasodilation to sodium nitroprusside was not different between young and old, male and female cerebral arteries post low and high PP. *n*= 6-11/group. A two-way ANOVA with Tukey's multiple comparisons was used. Data are mean ± SEM.
